# PI3Kγ pathway contributes to neuroinflammation and neuronal death induced by Zika virus infection

**DOI:** 10.1101/2024.12.20.629712

**Authors:** Danielle Cunha Teixeira, Gabriel Campolina-Silva, Fernanda Martins Marim, Felipe Rocha da Silva Santos, Celso Martins Queiroz-Junior, Pedro Augusto Carvalho Costa, Fernanda de Lima Tana, Luiz Pedro Costa-Souza, Jordane Clarisse Pimenta, Júlia Gomes Carvalho, Vinícius Amorim Beltrami, Felipe Emanuel Oliveira Rocha, Drielle Viana Vieira, Evandro Gonçalves Dornelas, Giovana Cougo Ferreira, Felipe Ferraz Dias, Pedro Pires Goulart Guimarães, Vinícius Toledo Ribas, Antonio Lucio Teixeira, Aline Silva de Miranda, Mauro Martins Teixeira, Daniele da Glória Souza, Vivian Vasconcelos Costa

**Affiliations:** Departament of Morphology; Instituto de Ciências Biológicas, Universidade Federal de Minas Gerais, Belo Horizonte, MG, Brazil; Departament of Biochemistry and Immunology; Instituto de Ciências Biológicas, Universidade Federal de Minas Gerais, Belo Horizonte, MG, Brazil; Departament of Microbiology; Instituto de Ciências Biológicas, Universidade Federal de Minas Gerais, Belo Horizonte, MG, Brazil; Department of Obstetrics, Gynecology and Reproduction, Faculty of Medicine, Université Laval, Quebec, QC, Canada; CHU de Quebec Research Center-Université Laval, Quebec, QC, Canada; Biggs Institute, University of Texas Health Science Center at San Antonio; Department of Biological Sciences, Universidade do Estado de Minas Gerais, MG, Brazil; Department of Physiology and Biophysics, Instituto de Ciências Biológicas, Universidade Federal de Minas Gerais, Belo Horizonte, MG, Brazil

**Keywords:** Zika virus (ZIKV), Neuroinflammation, Neuronal cell death, PI3K/AKT pathway, Phosphatidylinositol 3-kinase γ (PI3Kγ), Microglia activation

## Abstract

Zika virus (ZIKV) is an emerging arbovirus belonging to the Flaviviridae family and Orthoflavivirus genus, with a pronounced tropism for the central nervous system (CNS), where it induces neuroinflammation and neuronal death. ZIKV is known to exploit host cellular mechanisms, including the activation of survival pathways such as the PI3K/AKT signaling cascade, to evade apoptosis and enhance its replication. The phosphatidylinositol 3-kinase γ (PI3Kγ) pathway regulates critical cellular processes, including differentiation, recruitment, and survival, and is abundantly expressed in both brain tissue and leukocytes. This study aimed to investigate the role of the PI3Kγ pathway during ZIKV infection. Primary neuronal cultures from PI3Kγ-deficient mice (PI3Kγkd/kd) and human neuroblastoma SH-SY5Y cells treated with the PI3Kγ inhibitor AS605240 were infected with ZIKV to assess the impact of PI3Kγ signaling on viral replication and neuronal survival. Additionally, interferon α/β receptor knockout (A129) mice were treated with AS605240 either before or after ZIKV infection to evaluate the pathway’s role in neuroinflammation. In vitro, both genetic ablation and pharmacological inhibition of PI3Kγ suppressed ZIKV replication and prevented neuronal death. In vivo, mice treated with the PI3Kγ inhibitor exhibited enhanced protection against ZIKV infection, characterized by reduced viral load, and diminished brain and optic nerve damage. This neuroprotective effect correlated with altered astrocyte and microglia activation, marked by reduced TNF production in microglia. Furthermore, inhibition of PI3Kγ curtailed the recruitment and activation of CD8+ T cells and decreased the production of pro-inflammatory mediators, including IFN-γ and IL-17, in the brains of ZIKV-infected mice. These findings suggest that PI3Kγ activation facilitates ZIKV infection and exacerbates neuroinflammation. Pharmacological inhibition of the PI3Kγ pathway may offer therapeutic benefits by limiting viral replication and alleviating neuroinflammatory responses during ZIKV infection.

## INTRODUCTION

Zika virus (ZIKV) is a positive-sense single-stranded RNA arbovirus of the Flaviviridae family and Orthoflavivirus genus (Cunha et al., 2016). First isolated in 1947 from a rhesus monkey in Uganda, ZIKV later caused sporadic human infections in Africa and Asia before rapidly spreading to the Americas, including Brazil in 2015 (Dick, Kitchen, & Haddow, 1952). In that year, the World Health Organization declared ZIKV a public health emergency of international concern (Araujo, Ferreira, & Nascimento, 2016; Gulland, 2016; Lowe et al., 2018). It is hypothesized that mutations in the viral genome enhanced its adaptability and infectivity, facilitating its swift dissemination in the Americas and potentially contributing to worsened disease outcomes (Shan et al., 2020).

ZIKV infection has been linked to severe neurological outcomes, including microcephaly in neonates and Guillain-Barré syndrome (GBS) in adults (Araujo, Ferreira, & Nascimento, 2016; Gulland, 2016). Despite these associations, the underlying mechanisms of pathogenesis remain poorly understood. ZIKV exhibits tropism for the central nervous system (CNS), particularly targeting neural progenitor cells, microglia, and astrocytes, where it induces cell death, neuroinflammation, and neurodegeneration (Quincozes-Santos et al., 2023). In the CNS, ZIKV manipulates host signaling pathways to its advantage, including suppressing type I interferon (IFN-I) antiviral responses and activating the PI3K/AKT pathway to enhance cell survival and viral replication.

Therapies targeting host cell signaling pathways or enzymes involved in viral replication hold promise for treating ZIKV infection (Chan et al., 2018). In dengue virus (DENV) infection, treatment with the selective PI3Kγ inhibitor AS605240 in A129 mice reduced mortality, mitigated thrombocytopenia, and decreased vascular permeability and liver damage (Santos et al., 2024). Host cell kinases are increasingly recognized as key players in viral pathogenesis, as viruses exploit these enzymes to facilitate replication (Chu & Yang, 2007; Linero & Scolaro, 2009).

Phosphatidylinositol 3-kinase γ (PI3Kγ), a class I PI3K enzyme, is expressed in various cell types, including leukocytes and neurons. It regulates serine-threonine protein kinase (AKT) activation, which controls cell migration, growth, survival, autophagy, and apoptosis (Cantley & Neel, 1999; Hawkins & Stephens, 2015). Studies have shown that flaviviruses such as DENV and Japanese encephalitis virus activate the PI3K/AKT pathway to inhibit apoptosis and enhance viral replication (Lee, Liao, & Lin, 2005). Additionally, AKT phosphorylates ZIKV NS5 proteins, modulating RNA polymerase activity and promoting replication (Albentosa-González et al., 2021). Inhibition of the PI3K/AKT pathway using AR-12 has been shown to suppress ZIKV replication in neuronal cells and in A129 mice (Lee, Liao, & Lin, 2005). These findings highlight the PI3K/AKT pathway as a potential therapeutic target for ZIKV.

In this study, we investigated whether PI3Kγ blockade could prevent or mitigate neuroinflammation and neurodegeneration associated with ZIKV infection using both in vitro and in vivo models. Our results demonstrated that PI3Kγ inhibition reduces viral titers, prevents neuronal death, and suppresses the activation of inflammation-associated cells in the brains of ZIKV-infected mice. These findings suggest that targeting the PI3Kγ pathway may offer a novel therapeutic strategy to combat ZIKV-induced neuroinflammation and neurodegeneration.

## MATERIAL AND METHODS

This investigation adhered to the ethical and animal experiment regulations set forth by the Brazilian Government (law 11794/2008) and received approval of the Ethics Committee on Animal Experimentation at Universidade Federal de Minas Gerais (UFMG) (protocol n°. 72/2020). All experiments involving the ZIKV were conducted in strict accordance with biosafety level 2 regulations.

### Virus propagation and titration

A ZIKV isolate HS-2015-BA-01 was utilized in this study. It is a contemporary ZIKV isolate from 2015, during the Brazilian outbreak, and its genome is publicly available in GenBank under accession number KX520666.1. The virus was propagated in C6/36 *Aedes albopictus* cells and titrated via plaque assay, following established protocols (Costa *et al.,* 2014). For virus titration, virus-containing samples (cell culture supernatant, plasma, or mouse tissue extracts) were diluted and inoculated onto a monolayer of Vero E6 cells in an adsorption period of 1 hour and then incubated for 4 days, as previously described (Costa *et al.,* 2012). Results were expressed as Log_10_ plaque forming units (PFU) per mL or gram of tissue.

### Primary neuron culture system

Neuronal cultures were generated from embryonic brains of C57BL/6 wild-type and PI3Kγ^kd/kd^ mice, obtained at the 15^th^ day of gestation. Following enzymatic and mechanical dissociation (Alpuche-Lazcano *et al*., 2021), 3×10^4^ brain cells were plated in poly-L-ornithine-treated dishes and cultured in Neurobasal medium (Thermo Fisher Scientific), supplemented with N2 and B27, 2mM GlutaMAX (Thermo Fisher Scientific), 50 μg/mL streptomycin, and 50 U/mL penicillin (Gibco). After 5 days of culture in a humidified incubator (37°C, 5% CO2), primary neurons were inoculated with ZIKV (MOI of 0.1) for 1 hour, followed by an additional 48 h of culture in complete Neurobasal medium. Cell supernatant was collected for virus titration and ELISA, while cell damage/death was assessed through the LIVE/DEAD fluorescent viability assay (Invitrogen, #L3224) in the cellular fraction, as outlined in our previous work (Alpuche-Lazcano *et al*., 2021).

### Human Neuroblastoma Culture and Cytotoxicity Assay

SH-SY5Y neuroblastoma cells were obtained from the Rio de Janeiro Cell Bank (BCRJ) and cultured in Dulbecco’s Modified Eagle Medium F12 (DMEM F12) supplemented with 1% non-essential amino acids, 2 mM L-glutamine, 1 mM sodium pyruvate, and 10% fetal bovine serum. For the experiments, cells were seeded in 96-well microplates (1×10^5^ cells/well) and incubated at 37°C and 5% CO_2_ for 24 hours. Infection with ZIKV (MOI 1.0) occurred for 1 hour, followed by treatment with AS605240 at varying concentrations in DMEM medium supplemented with 5% serum. After 48 hours, the culture medium was collected for viral titration, LDH cell death assay (catalog no. K014, Bioclin, Belo Horizonte, Brazil), and ELISA. In addition to LDH, cell viability was assessed in the cellular fractions by MTT assay.

### Mouse strains

*In vivo* experiments were carried out using wild-type (WT) C57BL/6 mice (6-8 weeks old and 20-22g), obtained from the UFMG Specific-pathogen-free (SPF) Animal House, Brazil. Mice deficient for the catalytic portion of PI3KC (PI3KC^kd/kd^) on C57BL/6 background were originally purchased from The Jackson Laboratories (USA) and type I interferon receptor (IFN-α/βR^−/−^, referred to as A129^−/-^) knockout mice from SV129 background were originally acquired from B&K Universal Limited (United Kingdom) and maintained in the Animal Facility in Institute of Biological Science/UFMG. Mice were maintained under specific-pathogen-free conditions at 23°C on a 12-h light/12-h dark cycle with food and water provided ad libitum.

A129^−/-^ mice were inoculated via the tail vein with 200 μl PBS loaded with 4×10^3^ PFU of ZIKV as described elsewhere (Marim *et al*., 2021). In certain experiments, mice received subcutaneous treatment with the selective PI3KC pathway inhibitor AS605240 at a dose of 30 mg/kg body weight, either in a pre-treatment regimen (administration starting 1 hour before infection and every 24 hours for four days) or a post-treatment regimen (administration on days 2, 3, and 4 after infection). Vehicle mice infected with ZIKV alone were treated with a 5% DMSO solution diluted in saline, following the same pre-treatment or post-treatment scheme. The weight of the mice was monitored daily, and the intraocular pressure was measured on days 3 and 5 after infection. Mice considered severely ill with ≥ 20% weight reduction were euthanized. On day 5 post-infection (peak of ZIKV infection), the mice were euthanized to collect the brain and optic nerve.

### Assessing the intraocular pressure (IOP)

Intraocular pressure (IOP) was assessed in A129^−/-^ animals on days 0, 3, and 5 post-ZIKV infection. IOP measurements were conducted utilizing an applanation tonometer (Tono-Pen Vet - Reichert Technologies, NY, USA), following established protocols (Foureaux *et al*., 2015).

### Histopathology

Brain tissues from mock controls and ZIKV-infected mice were fixed in 10% formalin, embedded in paraffin (FFPE samples), and subjected to histopathological analyses. FFPE sections (5 µm thickness) were stained with hematoxylin and eosin (H&E), following our established protocol [21]. Histological damage in the cerebral cortex and hippocampus was assessed using a scoring system ranging from 0 (no damage) to 4 (necrosis). Meningeal inflammation was graded on a 0- to 4-point scale. The final score, representing the sum of cerebral cortex, hippocampus, and meningeal inflammation scores, was calculated, with a maximum possible score of 12 points (Costa *et al*., 2017; Marim *et al*., 2021). For the optic nerve analysis, the tissue was fixed in 10% paraformaldehyde diluted in PBS, incubated in 30% sucrose (Sigma-Aldrich) for 24h. Longitudinal cryosections (14μm) were generated using a cryostat (Leica), collected on glass slides, and stored at -20 °C until further evaluation.

### Immunohistochemistry

Sections of the cerebral cortex (5 µm thickness) were used for the staining of microglia and astrocytes. Following heat induced antigen retrieval in citrate buffer pH 6.0 (for Iba-1) or EDTA buffer pH 8.0 (for S100-β), tissues sections were blocked with Protein Block (#ab64226, Abcam) for 5 minutes and incubated overnight at 4°C with primary antibodies: anti-IBA-1 rabbit (PA5-21274, Invitrogen; 1:150) or anti-S100β rabbit (ab41548, Abcam; 1:75). After several washes in TBS, sections were exposed to biotinylated anti-rabbit IgG antibodies for 1 h, followed by ABC-HRP signal amplification (VECTASTAIN Elite ABC-HRP kit, Vector Laboratories, catalog no. PK-6200). 3,3′- Diaminobenzidine (DAB) was used as a chromogen to unravel the immunostainings. Image acquisition and analysis were performed using an Olympus BX 41 microscope (Olympus). The images showcased in the article serve as representatives of the experiments. For optic nerve immunostaining, sections were permeabilized with 0.3% Triton X-100 and 0.05% Tween-20 in PBS for 15 min and incubated with the blocking solution 10% normal goat serum (Sigma-Aldrich) in PBS at room temperature. Then, the sections were incubated overnight at 4 °C with anti-ß-III-Tubulin (Tuj1, 1:10.000, Biolegend), GFAP (1:2000, Cell Signaling), neurofilament-H (1:2000, Biolegend) and IBA-1 (1:2000, DAKO) primaries antibodies diluted in the same blocking solution. Sections were incubated with secondary antibodies Alexa-488 anti-mouse and Alexa-594 anti-rabbit (both 1:1000, ThermoFisher Scientific) for 1h at RT. The sections were counterstained with 4,6-diamidino-2-phenylindole (DAPI) and mounted in Fluomount (both ThermoFisher Scientific).

### Quantification of cytokine and chemokine concentrations in the brain

A range of inflammatory and neuromodulator molecules, including cytokines (IL-1β, IL-6, IFN-γ), chemokine (CX3CL-1), and the neurotrophin BDNF, were quantified in brain extracts using the mouse DuoSet ELISA kits from R&D Systems. Brain extracts were prepared by homogenizing 100 mg of tissue in 1 mL PBS supplemented with 0.1% Tween 20, 10 mM EDTA, 20 mM aprotinin, 0.1 nM PMSF, and 0.1 mM Benzethonium chloride. After centrifugation (9.184g, 15 min, 4°C), the supernatant was collected and used in the ELISA assays, following the manufacturer’s instructions (R&D Systems).

### Flow cytometry

The flow cytometry analysis followed the methodology described by Zaidan and collaborators (2022), focusing on various immune cell subpopulations identified by specific molecular markers. Brain tissues were macerated, suspended in RPMI culture medium and filtered through 70 μm cell strainer for cell isolation. Dead cells were excluded using Live/Dead (Invitrogen) for further labeling of extracellular and intracellular antigens after fixation and permeabilization (FoxP3 staining buffer set, eBioscience) according to the manufacturer’s instructions. LSR-FORTESSA equipment was used on acquisition. Data were analyzed by excluding debris, removing doublets with a forward scatter area (FSC-A) versus forward scatter height (FSC-H) gate, to avoid interruptions in the of flux. Cells were gated in function of time versus FSC-A, and so combinations of fluorochromes were done. These include activated microglial cells (CD45int CD11b+F4/80+MHCII+), CD4+ T cells (CD4+/CD3+), CD8+ T cells (CD8+/CD3+), regulatory T cells (Foxp3+/CD4+/CD3+), NK cells (CD3−/NK+), macrophages (CD45+/Gr-1−/F4/80+), monocytes (CD45+/Gr-1+/F4/80+), and neutrophils (CD45+/Gr-1+/F4/80−). The assessment also includes CD206 labeling and the quantification of TNF, Il-10, IL-17, IFN-γ, and iNOS within the cell subsets. The antibodies are listed in Table 1. Data analysis was performed using FlowJo V10.4.11.

### Statistical analysis

Statistical significance was determined utilizing GraphPad Prism 9.1.2 software, and the sample size was determined through analysis with G*Power 3.1 Software, guided by data from prior publications. No significant differences were found between the vehicle mock and the mock treated with AS605240 inhibitor groups in the in *vivo* experiments. Therefore, we used only the vehicle mock as the control group for the statistical tests. The results were analyzed using appropriate statistical tests, as indicated in figure legends. Data are represented as mean with standard deviation (SD).

## RESULTS

### PI3Kγ inhibition reduces neuronal damage induced by ZIKV infection in SH-SY5Y cells

Human neuroblastoma cells (SH-SY5Y) were infected with ZIKV (1×10C PFU, MOI 1) and treated for 48 h with the PI3Kγ inhibitor AS605240 (0.5, 5, and 50 μM). Supernatants were collected to assess cell death and viral titers, showing a dose-dependent effect of the inhibitor.

ZIKV infection triggered neuronal death and replicated in these cells. Treatment with the PI3K inhibitor after infection significantly decreased virus replication (Figure 1B) and cell death (Figure 1C). To confirm the inhibition of the PI3Ky signaling pathway, we quantified pAKT expression after treatment with 0.5 μM AS605240. While ZIKV increased pAKT levels, the group infected by ZIKV and treated with AS605240 did not show this increase, demonstrating the effectiveness of the pharmacological inhibition (Figure 1D-E).

**Figure 1.**
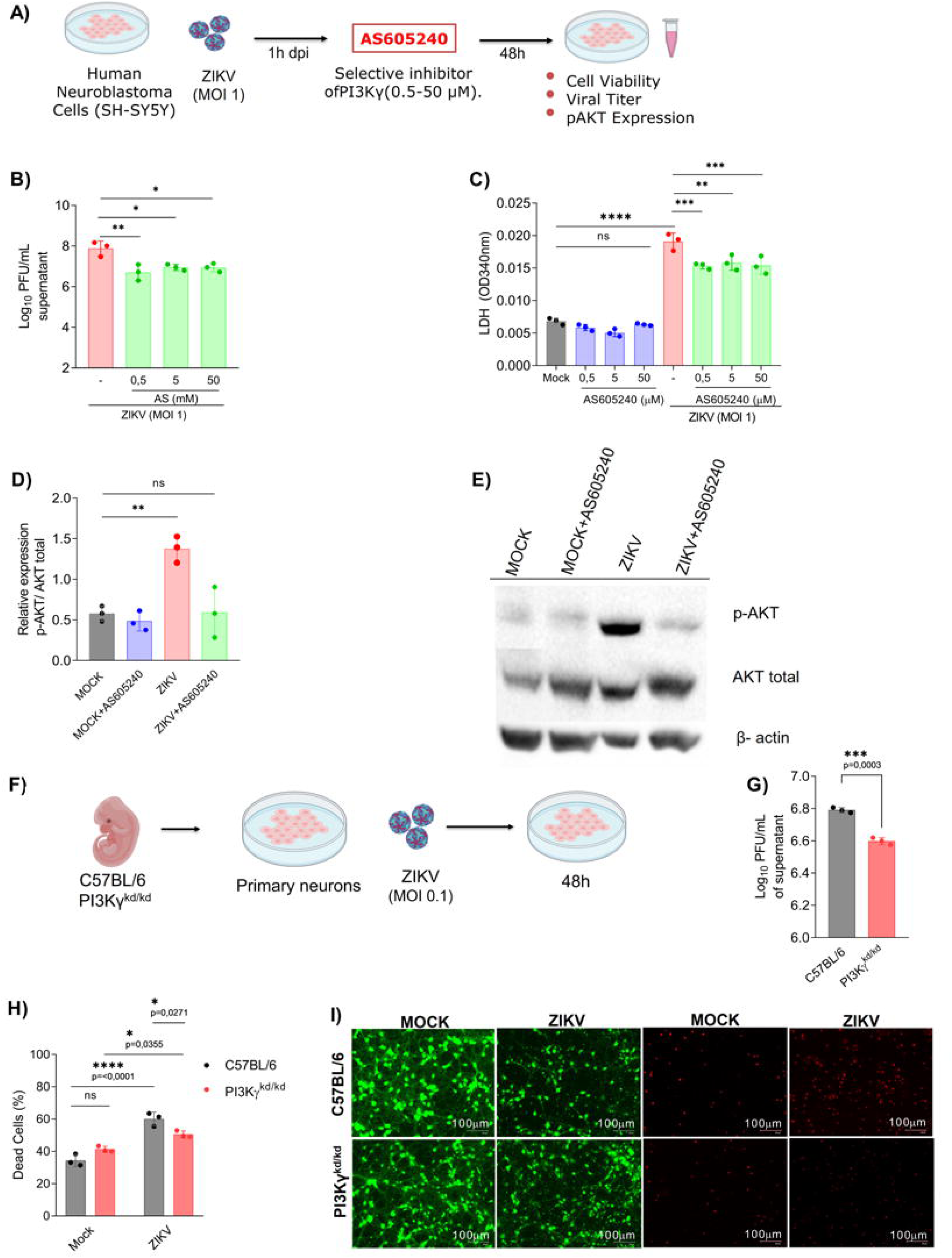
Reduction in neuronal death and viral titer in human neuroblastomas (SH-SY5Y) and in knockout mice for the PI3Ky enzyme. (A) Schematic representation of the experimental strategy to evaluate the therapeutic effects of the PI3Ky inhibitor AS605240 and pAKT expression. Firstly, SH-SY5Y was infected by ZIKV (MOI 1), then it was treated or not with the inhibitor AS605240 at concentrations of 0.5; 5; and 50 µM. After 48 hours, (B) viral titer and (C) cell death analyzes were performed in neuroblastoma cells. And to confirm the effect of the AS605240 inhibitor, we quantified the expression of pAKT, human neuroblastomas were infected by ZIKV (MOI 1) and subsequently treated with 0.5 μM AS605240. (D) Cells were used for pAKT quantification and (E) representative images. (F) Schematic representation of the experimental strategy, primary neuron culture was performed from WT mouse embryos; deficient for the catalytic subunit of the enzyme (PI3Kγkd/kd). After 48 hours of ZIKV infection, supernatant was collected to assess viral titer (G), cell death (H) was assessed using the LIVE/DEAD cell viability assay, and (I) representative images of infected and uninfected cells of neurons stained with calcein AM (green indicates live cells) and ethidium homodimer (red indicates dead cells). All results are expressed as mean and error bar indicates standard deviation (SD). Statistically significant differences were assessed by T-test (B), two-way ANOVA plus Tukey comparison test (C), one-way ANOVA plus Holm-Sidak, Sidak or Tukey comparison test (F-H). And were represented by *0.01 < P < 0.05; **0.001 < P < 0.01; ***P < 0.001; and ****P < 0.0001.

To assess whether this effect was indeed associated with the catalytic activity of PI3Kγ, we challenged neuronal cultures derived from embryos carrying a PI3Kγ loss-of-function mutation (PI3Kγ^kd/kd^), with ZIKV (MOI 0.1) for 48 hours (Figure 1F). Neurons lacking the catalytic activity of the PI3Kγ enzyme exhibited reduced viral titer (Figure 1G) and neuronal death (Figures 1H-I), compared to neurons established from wild-type embryos. Altogether, these results show that inhibition of the PI3Ky pathway is beneficial or protective during ZIKV infection *in vitro* in both human and murine neural cells.

### ZIKV infection drives AKT activation and PI3Ky Inhibition ameliorates disease outcomes in A129 Mice

Next, we investigated the influence of PI3Ky pathway during ZIKV infection *in vivo*. A129^−/-^ mice, which are susceptible to ZIKV infection (Costa *et al*., 2017), were intravenously inoculated with ZIKV (4×10^3^ PFU) and had their brain harvested at various time points post-infection (Figure 2A). ZIKV infection did not modulate PI3K-mediated AKT activation in the brain, as there was no change in pAKT levels (Figure 2B and C).

**Figure 2.**
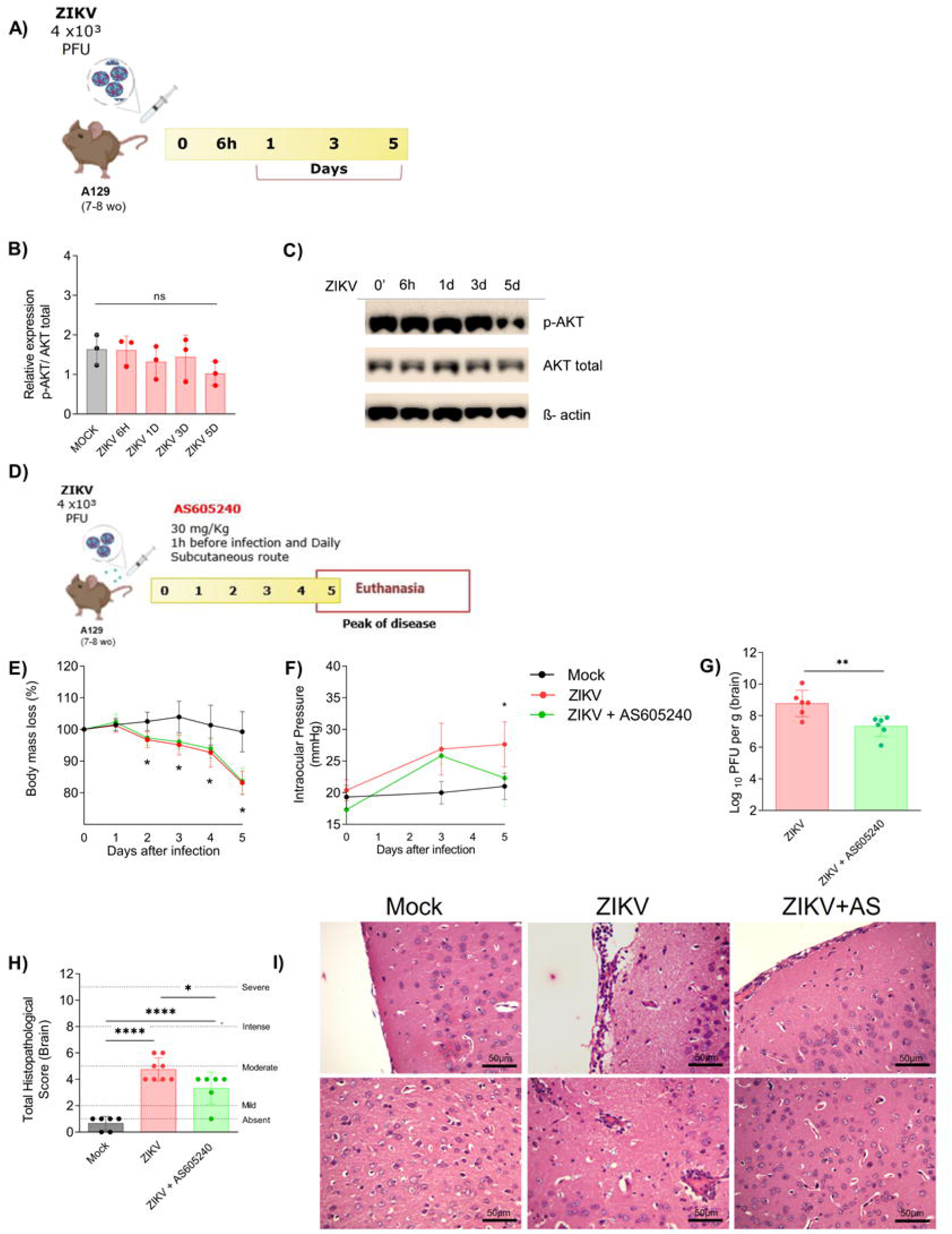
The PI3Ky pathway inhibitor improves clinical parameters, reduces viral titers and brain lesions in A129 mice infected by ZIKV. (A) Schematic representation of the experimental strategy, A129 mice were inoculated (iv) with 4×10^3^ PFU/200μL of ZIKV and then euthanized at different times of infection to evaluate pAKT expression. (B) quantification of pAKT expression in the animals’ brains after infection; (C) representative WB image. (D) Schematic representation of the experimental strategy to evaluate the role of the AS605240 inhibitor during ZIKV infection. AS605240 was administered 1 hour before infection and every 24 hours until euthanasia (day 5 of infection). (E) Body weight was assessed daily and (F) intraocular pressure was analyzed on days 0, 3 and 5. On day 5 of infection, the brain was harvested for (G) plaque assay analysis and for (H) semiquantitative analysis (histopathological scoring) after H&E staining of cerebral cortex and hippocampus sections. (I) Representative images of cerebral cortex sections. All results are expressed as mean and error bar indicates standard deviation (SD). Statistically significant differences were assessed by one-way ANOVA plus Sidak comparison test (B and H). two-way ANOVA plus Tukey comparison test (E,F) T-test (G), one-way ANOVA plus Sidak comparison test (H). And were represented by *0.01 < P < 0.05; **0.001 < P < 0.01; ***P < 0.001; and ****P < 0.0001.

Subsequently, we treated A129^−/-^ mice with a selective inhibitor of PI3Ky (AS605240; 30mg/kg), one hour before ZIKV infection and every 24 hours until euthanasia (Figure 2D). There was no difference in *in vivo* experiments between the mock receiving either vehicle or AS605240. Therefore, we used only the vehicle mock as the control group for the statistical analysis. ZIKV infection induced significant weight loss in mice, which was not reversed by the treatment with AS605240 (Figure 2E). However, inhibition of PI3Ky showed significant effects over the course of infection, by decreasing the IOP and ZIKV viral titer in the brain (Figure 2F and G). Moreover, while vehicle-treated ZIKV-infected mice exhibited marked meningitis, activated blood vessels in the cortex with signs of hyperemia and neighboring leukocytes, and indication of gliosis and necrosis, treatment of the mice with AS605240 attenuated the changes caused by ZIKV at 5 dpi (Figures 2H and I). Thus, the pharmacological inhibition of PI3Ky plays neuroprotective effects upon ZIKV challenge.

### PI3Ky inhibition reduces ZIKV-induced neuroinflammation without altering cytokine levels

It has been shown that ZIKV promotes neuroinflammation by infecting and activating glial cells, particularly microglia and astrocytes, leading to the release of pro-inflammatory cytokines (Marim *et al*., 2021). In the current experimental model, ZIKV infection induced a significant increase in the number of IBA-1^+^ macrophages/microglia and astrocytes (S100β^+^) in the brain cortex. While PI3Ky inhibition with AS605240 prevented changes in the number of these cells upon ZIKV infection (Figures 3A-C), it did not change the concentration of the pro-inflammatory cytokines IL-1β, IL-6, and IFN-γ or CX3CL-1, which were increased in the brain of infected mice (Figures 3D-H). Conversely, PI3Ky inhibition led to a significant increase of BDNF levels in the brain (Figure 3H), which may confer additional protective effects for neurons during infection.

**Figure 3.**
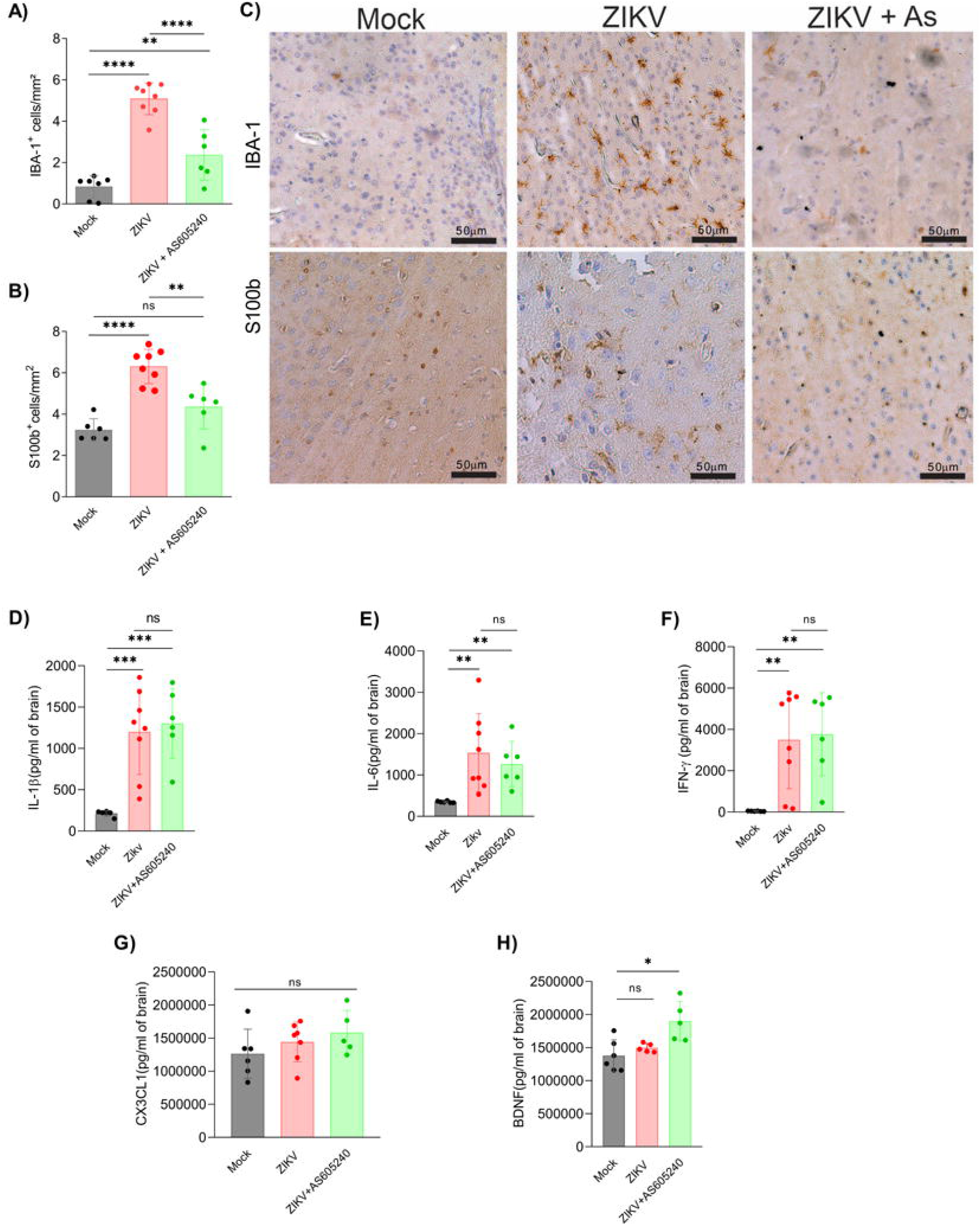
Treatment with a PI3Ky inhibitor reduced the activation of inflammatory cells in the brain induced by ZIKV infection without altering pro-inflammatory cytokines. Immunostaining of (A) IBA-1+, (B) S100-β was performed on mouse brain at 5dpi. (C) Representative images of cerebral cortex sections. (D–H) The production of chemokines/cytokines and neurotrophic factors were evaluated in mouse brains. All results are expressed as mean and error bar indicates standard deviation (SD). Statistically significant differences were assessed by one-way ANOVA plus Tukey comparison test. And were represented by *0.01 < P < 0.05; **0.001 < P < 0.01; ***P < 0.001; and ****P < 0.0001.

### PI3K inhibitor suppresses cell activation induced by ZIKV infection

The next step in our study was to investigate the effects of PI3K inhibition in the brain cell populations of ZIKV-infected mice. For this study, A129^−/-^ mice were intravenously inoculated with 4×10³ PFU of ZIKV. One hour prior to infection, the mice were treated with AS605240 (30 mg/kg), with subsequent doses administered every 24 hours until 5 days post-infection (dpi), when the brains were collected for further analysis using flow cytometry (Figure 4A). Representative density plots show the gating strategy employed for activated microglial cells (CD45intCD11b+F4/80+MHCII+) (Figure 4B).

**Figure 4.**
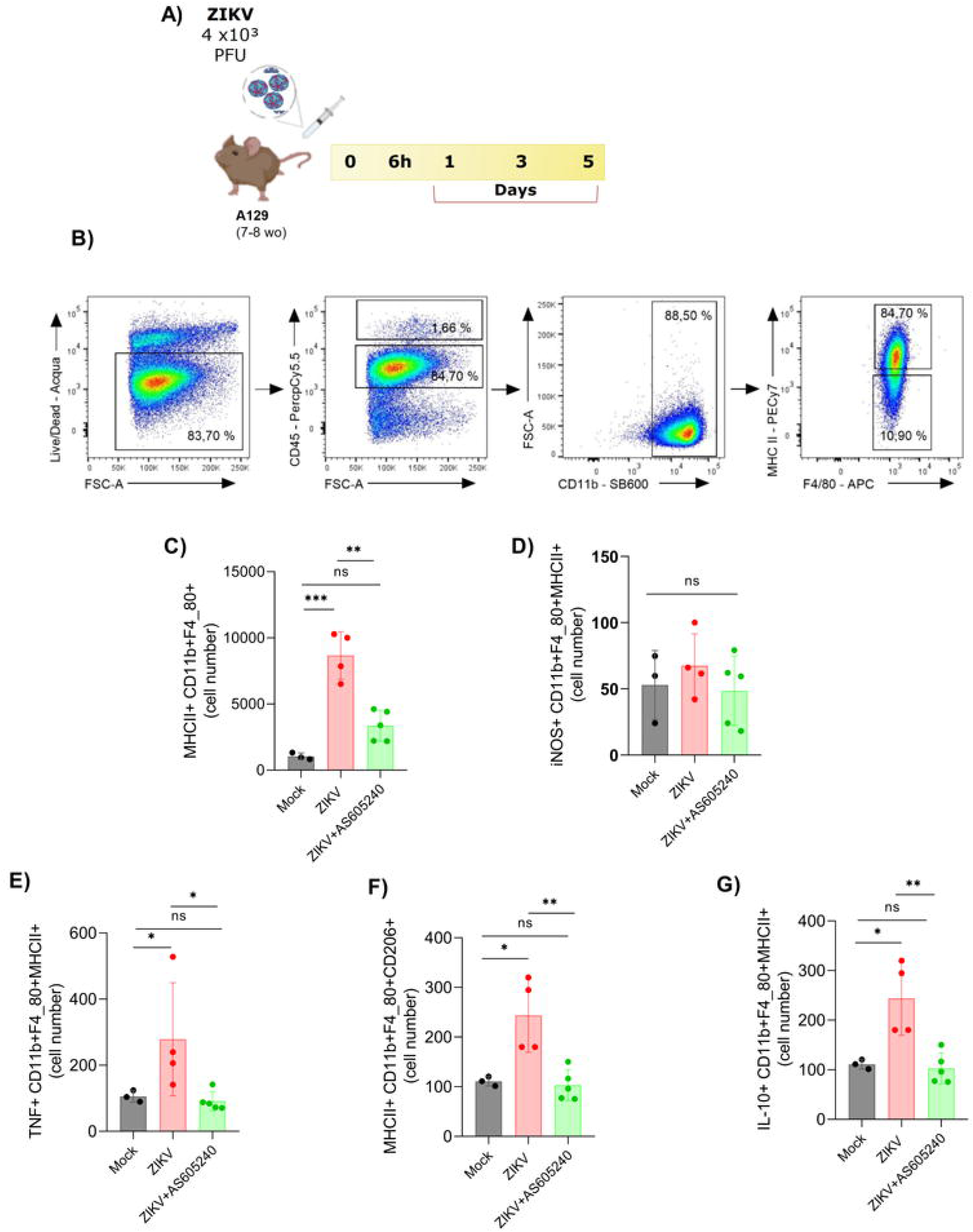
The PI3Ky pathway inhibitor decreases microglial activation and TNF production triggered by ZIKV infection. (A) Scheme of infection with ZIKV and treatment with AS605240. (B) Representative density plots of the proportion of (CD45^inter^), (CD11b+), activated microglia (F4/80+/MHC-I+) from a single ZIKV-infected brain. (C) Activated microglia (CD45intCD11b+F4/80+MHC-1+). (D) Activated microglia expressing mannose receptor (CD45intCD11b+F4/80+MHC-1+CD206+). (E) IL10-producing microglia (CD45intCD11b+F4/80+MHC-1+IL-10+), (F) iNos activity (CD45intCD11b+F4/80+MHC-1+iNos+) and (G) TNF release (CD45intCD11b+F4/80+MHC-1+TNF+). All results are expressed as mean and error bar indicates standard deviation (SD). Statistically significant differences were assessed by one-way ANOVA plus Sidak’s (C, E,F) or Tukey’s (D, G) comparison test. And were represented by *0.01 < P < 0.05; **0.001 < P < 0.01; ***P < 0.001; and ****P < 0.0001.

ZIKV infection induced the activation of microglia and the release of both anti-inflammatory (IL-10) and pro-inflammatory (TNF) mediators (Figure 4C-G). Conversely, treatment with AS605240 prevented the activation of microglia induced by ZIKV infection (Figure 4C). To better determine the inflammatory profile, we characterized the expression of iNOS and various cytokines in these cells. While AS605240 treatment did not alter the number of microglial cells expressing iNOS (Figure 4D), it significantly reduced the production of the pro-inflammatory cytokine TNF (Figure 4E) in activated microglia during ZIKV infection. Additionally, we examined the expression of the mannose receptor (CD206+) and observed that AS605240 treatment not only decreased CD206 receptor expression but also reduced IL-10 production (Figure 4F-G) in activated microglia compared to the ZIKV group. No changes were observed in the number or cytokine production by monocytes, macrophages, neutrophils, and dendritic cells in the AS605240-treated group compared to the mock and ZIKV groups (Figure S1A-O). Overall, these findings suggest that AS605240 treatment specifically prevents microglial activation and modulates the production of pro- and anti-inflammatory cytokines, potentially ameliorating ZIKV infection.

We also investigated the role of PI3Kγ inhibitor in CD8+ T cells in the context of ZIKV infection. Representative density plots show the control strategy used for CD8+ T cells (CD45^high^CD3+CD8+) (Figure 5A). ZIKV induced an increase in the number of CD8+ T cells while the treatment with AS605240 inhibitor reduced these numbers (Figure 5B). Concomitantly, ZIKV infection increased the release of IFN-γ and IL-17 by TCD8+ cells, but treatment with AS605240 inhibitor reduced the production of these mediators (Figure 5C-D). No changes in the number or cytokine production by CD4+ T cells, Treg+ cells, NK+ cells, NKT+ cells were observed in the ZIKV vehicle group compared to the ZIKV-infected and AS605240-treated group (Figure S2A-L). Our findings also suggest that targeting the PI3Ky pathway could potentially alleviate the effects of ZIKV infection by reducing CNS inflammation.

**Figure 5.**
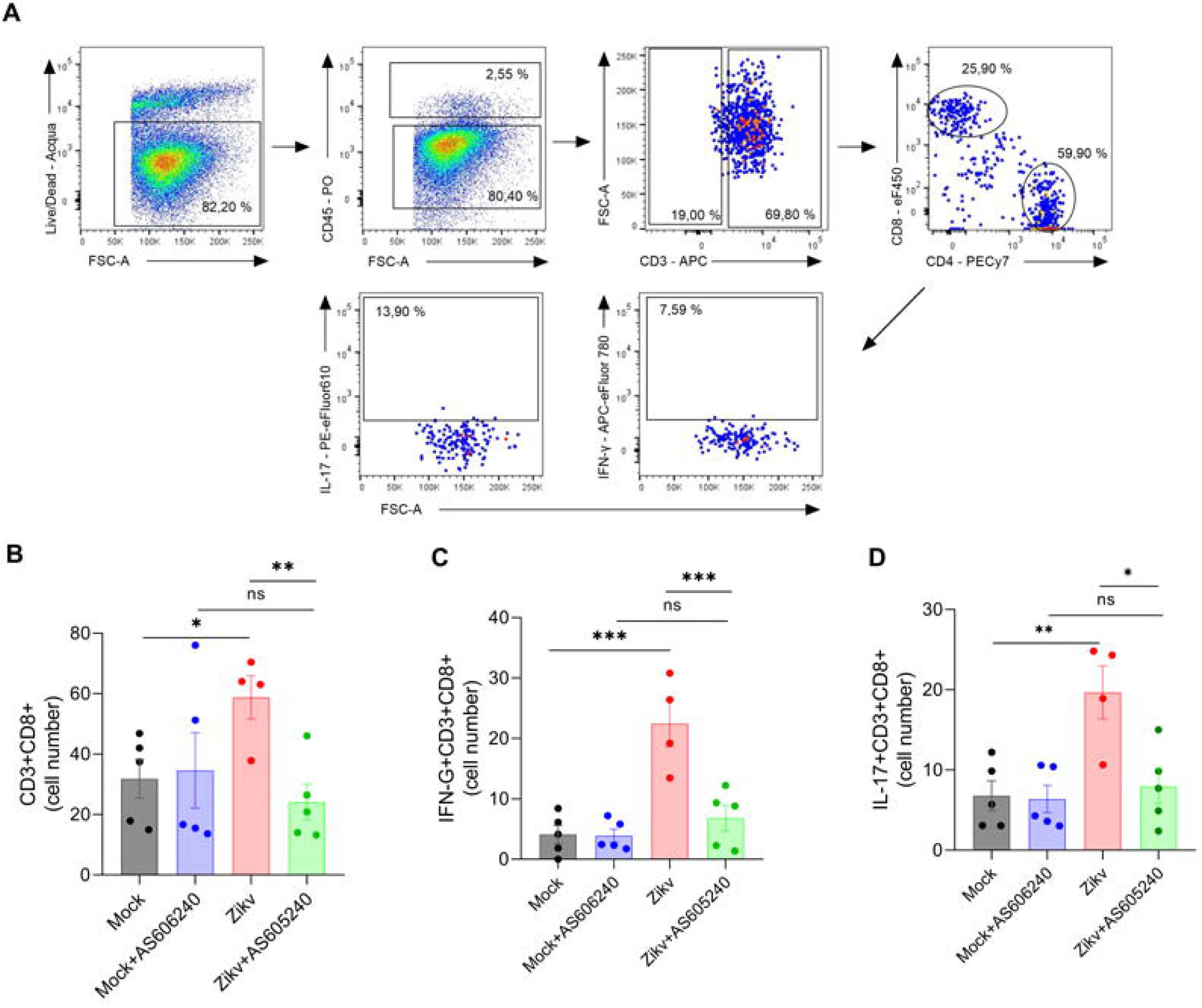
Treatment with a PI3Kγ pathway inhibitor decreases the activation of CD8 T and the production of IFN- and IL-17 triggered by ZIKV infection. (A) Representative density plots of the proportion of (CD45^high^), (CD3+), CD8 T cells (CD3+CD8+) from a single ZIKV-infected brain. (B) CD8 T cell (CD3+CD8+). (C) IFN-C-producing CD8 T cell expressing (CD45+CD3+CD8+IFN-C+), (E) IL17-producing CD8 T cell (CD45+CD3+CD80+IL-17+). All results are expressed as mean and error bar indicates standard deviation (SD). Statistically significant differences were assessed by one-way ANOVA plus Tukey comparison test. And were represented by *0.01 < P < 0.05; **0.001 < P < 0.01; ***P < 0.001; and ****P < 0.0001.

### Therapeutic inhibition of PI3Ky reduces ZIKA replication and mitigates brain and optic nerve damage

To evaluate the therapeutic potential of PI3Ky inhibition, AS605240 treatment (30mg/kg) was initiated on the 2^nd^ day post-infection and continued daily until euthanasia (Figure 6A). Brain, optic nerve, and eye samples were collected five days post-infection. Significant body weight loss occurred due to ZIKV infection, but AS605240 treatment initiated on the 2^nd^ day post-infection did not influence this effect (Figure 6B). ZIKV infection led to increased intraocular pressure (IOP), with no significant differences observed compared to the infected and treated group (Figure 6C). Nevertheless, AS605240 treatment effectively reduced viral loads (Figure 6D), inflammation, and histopathological damages in the brains of treated mice compared to those in the ZIKV-vehicle group (Figures 6E-F). Immunostaining analysis revealed a decrease in IBA-1+ cells without changes in S100β+ cells in treated mice compared to the ZIKV-vehicle group (Figures 6G-I). In the optic nerve, ZIKV infection led to axonal damage showed by fragmentation of ß-III-Tubulin (Tuj1) and neurofilament H immunostainings (Figures 7A-B). However, AS605240 treatment reduced the fragmentation of ß-III-Tubulin (Tuj1) and neurofilament H staining, suggesting that it promoted axonal protection (Figures 7A-B). There were no obvious changes in the GFAP+ cells in the optic nerve (Figure 7A). In uninfected mice, IBA-1+ macrophages/microglia cells showed normal morphology with thin process (Figure 7B). ZIKV infection induced changes in the morphology of IBA-1+ cells, which showed thicker processes. PI3Ky inhibition with AS605240 did not prevent these changes induced by ZIKV infection (Figure 7B). Thus, PI3Ky inhibition presents promising therapeutic opportunities for mitigating ZIKV replication and associated central nervous system injuries.

**Figure 6.**
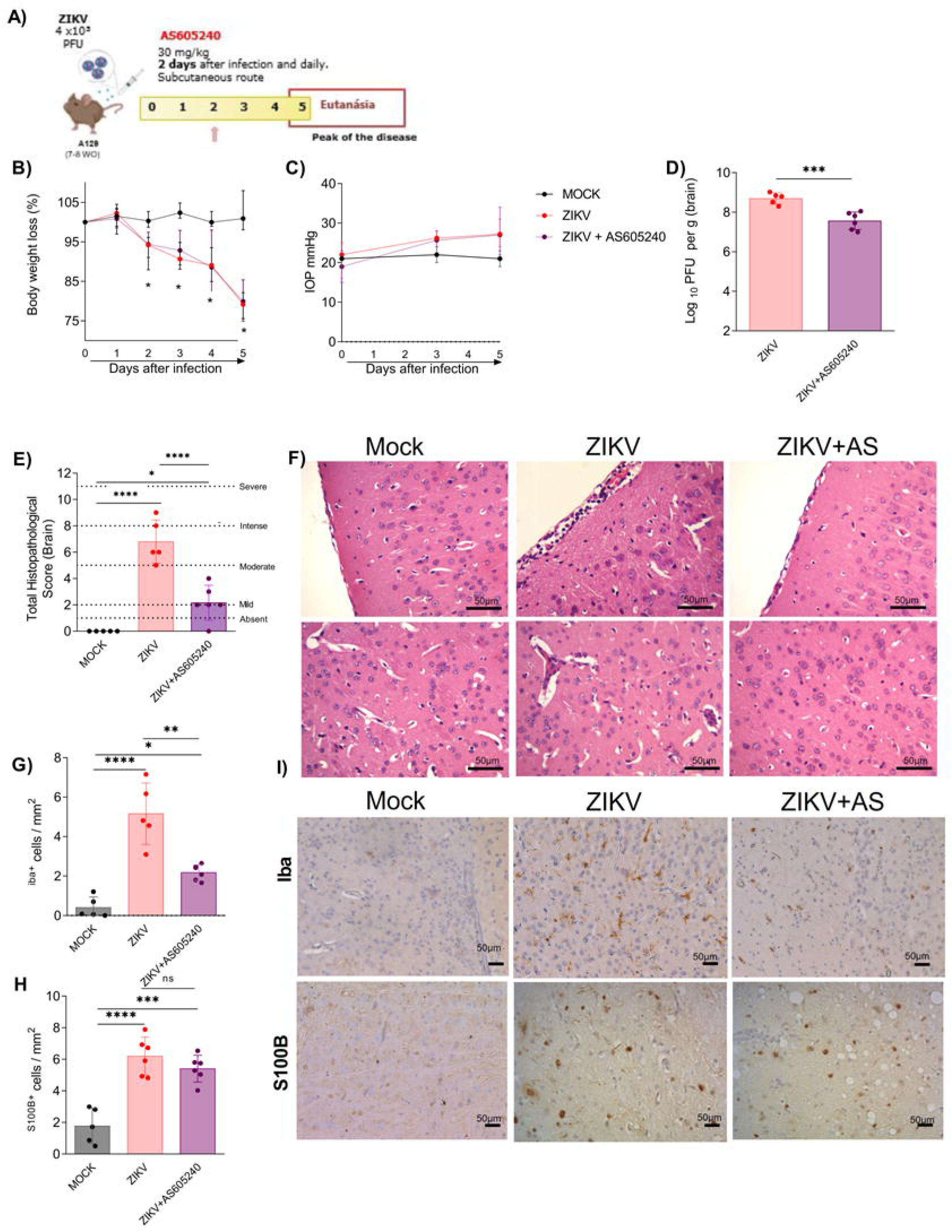
Therapeutic administration with the PI3Ky pathway inhibitor reduces viral titers, injury and inflammation induced by ZIKV. (A) Schematic representation of the experimental strategy, A129 mice were inoculated (iv) with 4×10^3^ PFU/200μL of ZIKV. 2 days after infection, mice were treated with AS605240 (30 mg/kg) every 24 hrs for 5 days. (B) Body weight was assessed daily and (C) intraocular pressure was analyzed on days 0, 3 and 5. On day 5 of infection, the brain was harvested for (D) plaque assay analysis and for (E) semiquantitative analysis (histopathological scoring) after H&E staining of cerebral cortex and hippocampus sections. (F) Representative images of cerebral cortex sections. Immunostainings of (G) IBA-1+, (H) S100-β, and (I) Representative images of cerebral cortex sections. All results are expressed as mean and error bar indicates standard deviation (SD). Statistically significant differences were assessed by two-way ANOVA plus Tukey comparison test (B,C), T-test (D), one-way ANOVA plus Sidak’s (E) orTukey’s (G,H) comparison test. And were represented by *0.01 < P < 0.05; **0.001 < P < 0.01; ***P < 0.001; and ****P < 0.0001.

**Figure 7.**
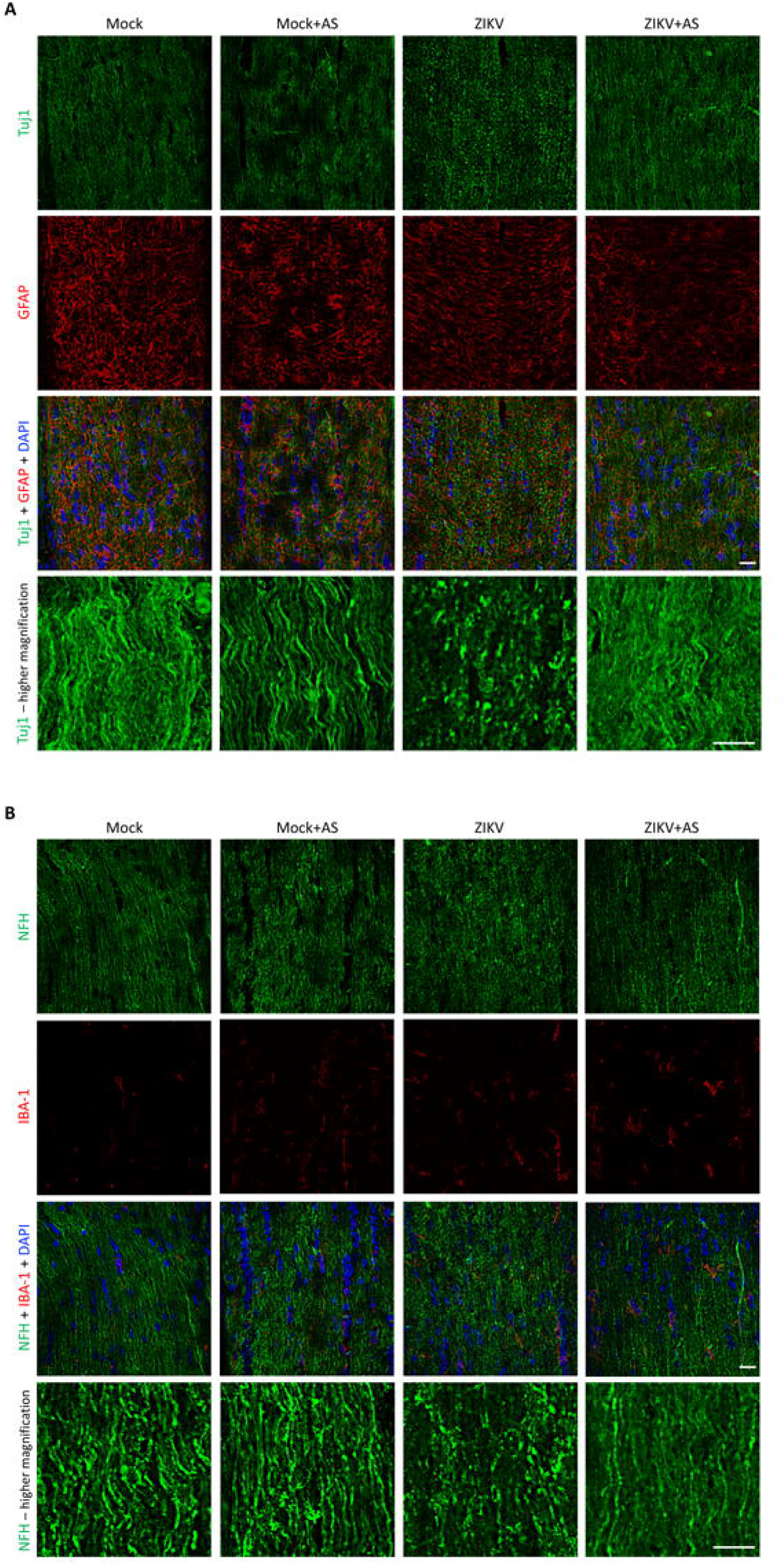
Therapeutic administration with the PI3Ky pathway inhibitor reduces axonal damage in the optic nerve induced by ZIKV. Representative images of optic nerve cryosections showing immunostainings for ß-III-Tubulin (Tuj1) and GFAP (A), neurofilament H (NFH) and IBA-1 (B). (A, B) Lower rows show higher magnification images of ß-III-Tubulin (Tuj1 - A) and neurofilament H (NFH - B) immunostainings. Scale bars: 20 µm and 10 µm (higher magnification).

## Discussion

After Zika outbreaks in the Americas, studies demonstrated that ZIKV is neurotropic and triggers an inflammatory response that compromises the integrity of the Blood-Brain Barrier (BBB), facilitating its spread to other CNS cells, such as neurons, astrocytes and microglia. In response to infection, the cells release pro-inflammatory cytokines and chemokines that contribute to imbalance in homeostasis and neuroinflammation (Clé *et al*., 2020). Because of the ZIKV-related neuronal cell death, the infection can cause different neurological disorders, such as microcephaly (Mlakar *et al*., 2016). Therefore, it is urgent to understand cellular mechanisms that the virus can use to benefit itself and develop therapies. In the current study, we investigated the potential neuroprotective effect of inhibiting the PI3Kγ pathway during ZIKV infection using a pharmacological strategy (the inhibitor AS605240) or a genetic approach (PI3Kγ kd/kd deficiente mice), both *in vitro* and *in vivo*. Our main findings were: (I) Reduction in neuronal death and viral titer in human neuroblastoma cells (SH-SY5Y) treated with AS605240 inhibitor or neurons derived from knockout embryos for the PI3Kγkd/kd catalytic enzyme; (II) Reduction in viral titers, injury, activation of inflammatory cells and release of pro-inflammatory cytokines in the brain of mice previously treated with AS605240 inhibitor and infected by ZIKV; (III) Reduction of viral titers, injury, activation of inflammatory cells in the brain and axonal damage in the optic nerve of mice infected by ZIKV and therapeutically treated with AS605240 inhibitor.

The PI3K/Akt pathway plays a crucial role in controlling essential cellular functions, including cell migration, survival, metabolism, and nutrition (Cantley; Neel, 1999; Hawkins; Stephens, 2015). The PI3Kγ isoform is especially expressed in leukocyte cells, endothelial cells, glial cells and neuronal cells (Lanahan; Wymann; Lucas, 2022). Due to the fundamental role of PI3Ky in leukocyte migration, this pathway has become the target of studies in inflammatory diseases, whether non-infectious or infectious (Bellozi *et al*., 2016; Rodrigues *et al*., 2010). Previous studies have highlighted the importance of the PI3Kγ pathway in various viral infections (Dunn; Connor, 2012). For example, the PI3K/AKT pathway regulates the replication of different viruses, including Japanese encephalitis virus (JEV) and DENV-2 viruses (Lee; Liao; Lin, 2005), hepatitis B and C viruses (He *et al*., 2002; Shih *et al*., 2000), Vaccinia and Cowpox viruses (Soares *et al*., 2009), Usutu virus (USUV), West Nile virus (WNV) (Albentosa-González *et al*., 2021), among others. These viruses influence the PI3K/AKT pathway in several ways, such as the entry of the virus into cells, the translation of the viral protein and mainly cell survival, as this mechanism postpones the apoptosis of the host cell and ensures the persistence of the virus during infection (Diehl; Schaal, 2013). When we investigated the selective protection of PI3Kγ with AS605240 in human neuroblastomas or in primary neurons derived from embryos with PI3Kγkd/kd, which are extremely important cells in ZIKV infection, we observed a significant reduction in viral replication and cell mortality when compared to the control group. These findings corroborate a previous study where researchers show that treatment with the celecoxib-derived kinase inhibitor AR-12 (OSU-03012) inhibited Zika virus through downregulation of PI3K/Akt in neuronal cells (Chan *et al*., 2018). Furthermore, it has been demonstrated that during ZIKV infection, host cell AKT interacts with the virus’s NS5 protein, contributing to viral replication. These results were confirmed by treatment with the specific AKT inhibitor, MK-2206, which impairs viral load in cells heavily infected with ZIKV (Albentosa-González *et al*., 2021).

We next investigated the effects of pharmacological inhibition of PI3Kγ on ZIKV *in vivo*. We found that the pharmacological inhibitor of the PI3Kγ pathway (AS605240), administered before or after ZIKV infection, impairs viral replication in the brain of animals. This finding corroborates existing literature, which indicates that AKT downregulation reduces viral replication in the brains of ZIKV-infected A129 mice (Chan *et al*., 2018). Furthermore, our results demonstrated that pharmacological inhibition of the PI3Kγ pathway also minimizes the lesions caused by viruses in the brains and optic nerve of infected mice.

Subsequently, we investigated potential ZIKV target cells that could contribute to increased inflammation in the brain, such as microglia and astrocytes (Lum *et al*., 2017; Veilleux; Eugenin, 2023). Our results confirmed that ZIKV triggered an increase in the number of IBA-1 and S100b cells. Microglia, when activated during ZIKV infection, produces high levels of neurotoxic and pro-inflammatory factors associated to neuroinflammation, favoring damaging and the spread of the virus in the brain (Lum *et al*., 2017). Similarly, astrocytes, when infected, contribute to viral replication, resulting in neuroinflammation and neuronal death (Veilleux; Eugenin, 2023). Treatment with a PI3K inhibitor in the group of mice infected by ZIKV was able to prevent both the increase in cells positive for IBA-1 and S100b when compared to the group infected with ZIKV but not treated. Accordingly, in the context of inflammatory stimuli, the PI3K/AKT pathway plays an important role in activated microglia, such as the activation of NF-κB-dependent inflammatory response pathways (Saponaro *et al*., 2012). Likewise, studies carried out by our group demonstrated that PI3Kγ deficient mice (PI3Kγ-/-) infected with malaria had a reduction in hemorrhage, vascular obstruction and accumulation of leukocytes, as well as less microglial activation (Iba-1 + reactive microglia) in the brain (Lacerda-Queiroz *et al*., 2015). It has also been described that there is a correlation between the positive regulation of the PI3K/AKT pathway and the reactivity of astrocytes (Cheng *et al*., 2020; Zhang *et al*., 2024). These reactive astrocytes exhibited increased levels of PI3K, pAKT and AKT against inflammatory stimuli caused by LPS+IFNγ (Cheng *et al*., 2020)

Although we observed that inhibition of PI3Ky prevents the increase in inflammatory cells in the brain of A129 mice, it is interesting to note that there is no difference in the levels of pro-inflammatory mediators or chemokines between the infected groups. This same phenotype was detected in other contexts. Although ZIKV-induced neurodegeneration was prevented by memantine treatment, the production of inflammatory mediators and weight loss in IFN-α/βR -/- mice were not affected (Costa *et al*., 2017). Likewise, a tryptophan inhibitor also reduced the number of microglia and astrocytes, although it did not interfere with ZIKV viral titers and inflammatory mediators (Marim *et al*., 2021). Interestingly, our results demonstrated that BDNF, a neurotrophic factor responsible for promoting the survival, maintenance and reorganization of the neuronal microenvironment in the CNS (Lima Giacobbo *et al*., 2019), levels are increased only in the group infected and treated with AS605240. This increase in BDNF may be associated with cellular mechanisms to ensure brain homeostasis and infection control, as the treatment contributed to the reduction of lesions and cellular activation. Whereas neuroinflammation is regulated by factors that are also involved in modulating BDNF expression and patients with neuropsychiatric and neurodegenerative disorders often present reduced concentrations of BDNF in the blood and brain (Diniz; Teixeira, 2011; Lima Giacobbo *et al*., 2019). Previous studies with ZIKV demonstrated an important decrease in BDNF levels in the placentas of ZIKV-infected patients, especially in groups of babies with microcephaly (Rabelo *et al*., 2020).

In a cellular context, ZIKV infection in A129 mice caused an increase in microglial activation, increased the production of the pro-inflammatory mediator TNF and, interestingly, also increased the expression of the CD206+ receptor and the production of the anti-inflammatory cytokine IL-10. When we evaluated mice treated with the inhibitor AS605240, the drug was still able to inhibit the increase in microglial activation along with the expression of TNF, CD206+ receptor and IL-10, confirming that inhibition of PI3Kγ may be beneficial in ZIKV infection. Furthermore, these findings suggest the possibility of a control of homeostasis, since it has already been demonstrated that IL-10 could modulate the PI3K pathway in microglia, resulting in the inhibition of the production and release of TNF-α induced by LPS (Lum *et al*., 2017). In line with our data, previous work described the PI3K/AKT pathway as modulated by activated microglia. Once activated, this pathway results in increased production of pro-inflammatory factors, such as TNF, and the acquisition of a neurotoxic phenotype (Saponaro *et al*., 2012).

It is known that the activation of microglia and astrocytes during a viral infection contributes to the influx of TCD8 effector cells into the CNS (Garber *et al*., 2019). CD8+ T Cells in Ifnar1^−/-^ mice during ZIKV infection are responsible for preventing viral replication through indirect mechanisms, such as the release of cytokines (Balint *et al*., 2024). Although CD8 + T cells have antiviral activity in the brain, limiting ZIKV infection of neurons, their activity also contributes to the stimulation of ZIKV-associated paralysis (Diniz; Teixeira, 2011). When analyzing the group previously treated with the inhibitor AS605240, we observed that the treatment prevented the increase in CD8+ T cells and, consequently, the increase in the mediators IFN-γ and IL-17. In West Nive Virus and ZIKV infection, excess of these pro-inflammatory cytokines, such as IFN-γ, mediated by CD8+ T cells, can damage neuronal structures and promote neuronal apoptosis (Garber *et al*., 2019; Kreutzfeldt *et al*., 2013; Satchidanandam, 2021). Furthermore, PI3Kγ has also been described to play a crucial role in the migration of effector CD8 T cells to inflammatory sites, demonstrating that p110γ-deficient effector CD8 T cells exhibit impaired or reduced migration both in vitro and in vivo (Martin *et al*., 2008). Thus, our results suggest that inhibition of PI3Kγ is important to control CD8 T cell migration, the neurotoxic environment and cell death provided by CD8 T cells during ZIKV infection.

## Conclusion

In summary, PI3Kγ inhibition exerted a neuroprotective effect by attenuating viral replication and CNS damage associated with ZIKV infection. This favorable outcome seems to stem from the decreased activation of specific cells, including microglia and CD8 T cells, leading to a subsequent reduction in the release of pro-inflammatory mediators by these cells. While PI3Kγ inhibition holds promise as a therapeutic strategy for treating ZIKV infection, further investigations are warranted to fully elucidate the underlying neuroprotective mechanisms.

## Acknowledgements

We are grateful to Ilma Marçal de Souza, Rosemeire Oliveira, Letícia Soldati and Tânia Colina for their technical assistance. The authors thank L’Oréal-UNESCO-ABC ‘Para Mulheres na Ciência’ prize granted to VVC.

