## Supplemental Table 1 for "PI3Kγ pathway contributes to neuroinflammation and neuronal death induced by Zika virus infection"

| Antigen | [Fluorochrome](http://www.fluorochrome.com/) |  | Clone | Concentration | Company |
| --- | --- | --- | --- | --- | --- |
| TNF-α | eFluor450 |  | MP6-XT22 | 1/1000 | ThermoFhisher |
| CD8a | eFluor 450 |  | 53-6.7 | 1/1300 | ThermoFhisher |
| LIVE/DEAD | Acqua |  |  | 1/1000 | ThermoFhisher |
| Streptavidin | Pacific Orange |  |  | 1/200 | ThermoFhisher |
| CD45 | Pacific Orange |  | 30-F11 | 1/200 | ThermoFhisher |
| CD11b | Super Bright 600 |  | M1/70 | 1/500 | ThermoFhisher |
| CD206 | PE |  | MR6F3 | 1/300 | ThermoFhisher |
| Foxp3 | PE |  | 150D/E4 | 1/100 | ThermoFhisher |
| IL-17 | PE-eFluor610 |  | eBio17B7 | 1/100 | ThermoFhisher |
| iNos | PE-eFluor610 |  | CXNFT | 1/500 | ThermoFhisher |
| CD45 | PerCP-Cy5.5 |  | Ly-5.2 | 1/200 | BioLegend |
| NK | PerCP-Cy5.5 |  | PK136 | 1/150 | ThermoFhisher |
| CD4 | PE-Cyanine7 |  | GK1.5 | 1/3000 | ThermoFhisher |
| F4/80 | APC |  | BM8 | 1/200 | ThermoFhisher |
| IL-10 | AlexaFluor 700 |  | JES5-16E3 | 1/100 | ThermoFhisher |
| MHC-II | APC-eFluor 780 |  | M5/114.15.2 | 1/500 | ThermoFhisher |
| CD3 | FITC |  | 17A2 | 1/400 | ThermoFhisher |
| IFN-γ | APC-eFluor 780 |  | XMG1.2 | 1/200 | ThermoFhisher |
| Gr-1 | Biotin |  | RB6-8C5 | 1/500 | Biolegend |
