## Supplementary figures and images for "PI3Kγ pathway contributes to neuroinflammation and neuronal death induced by Zika virus infection"

### Supplemental Figure 1

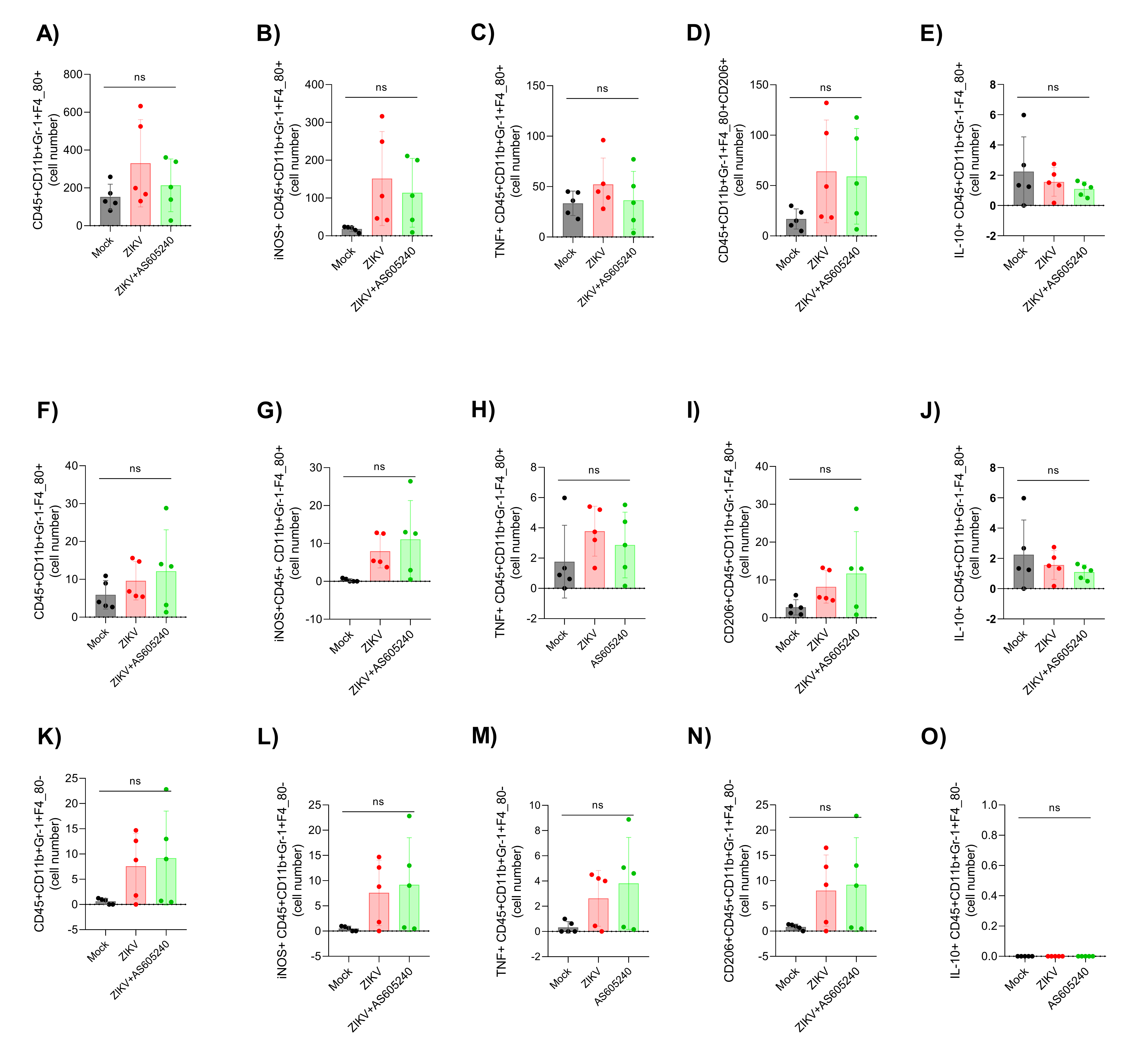

### Supplemental Figure 2

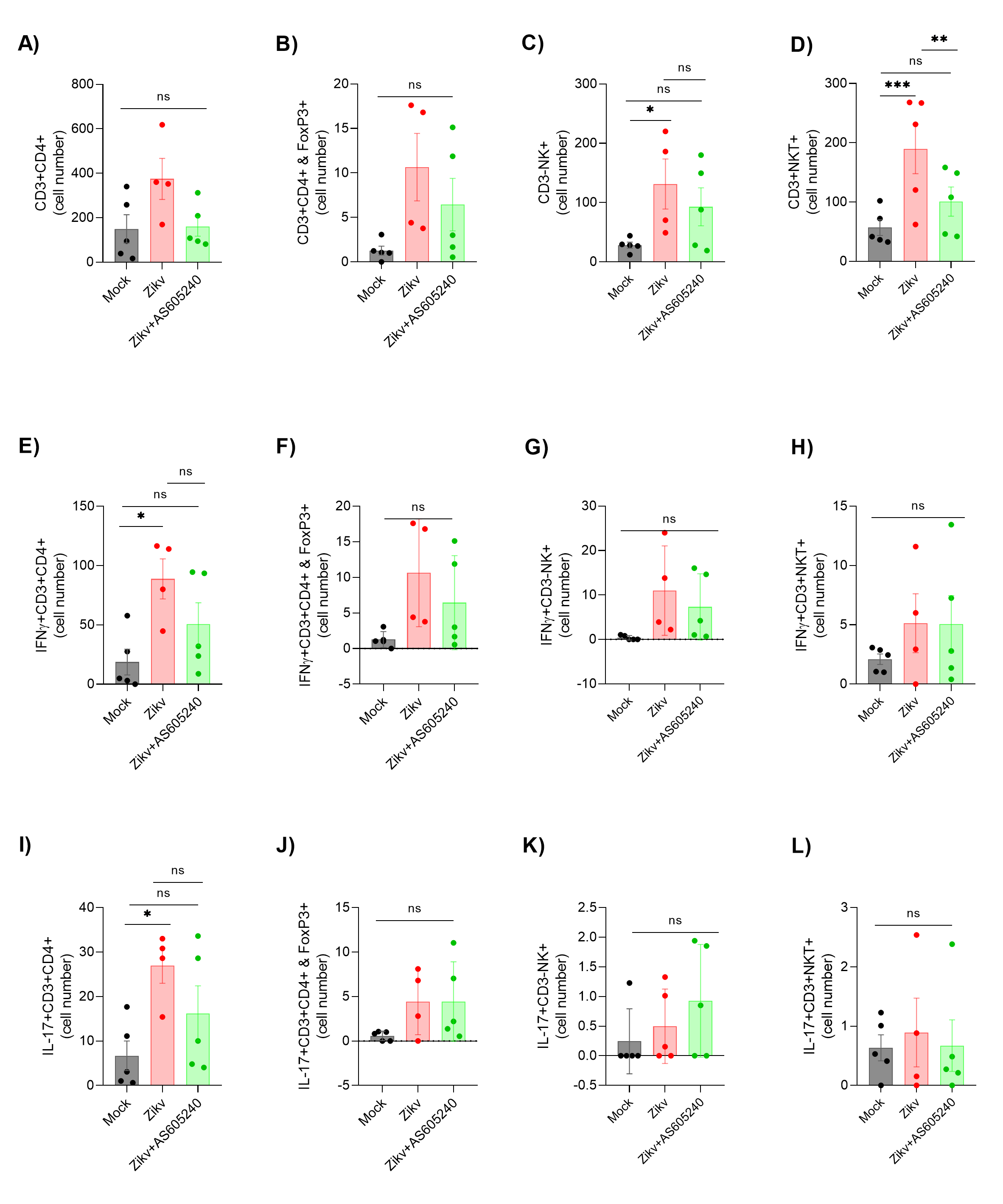
